# Brain Entropy: Linking Brain Structure to Task Activation

**DOI:** 10.1101/2025.10.27.684734

**Authors:** Donghui Song

## Abstract

Understanding how the brain’s complex and diverse functions emerge from its structure has long been a central goal of neuroscience. Entropy, a measure of system irregularity, has become a powerful tool for quantifying the dynamics of brain activity. While previous fMRI-based brain entropy (BEN) studies have established its relationships with brain morphology (e.g., gray matter volume, GMV) and task activation, the pathway from structure to function remains incompletely understood. Specifically, it is unclear whether controlling for brain morphology (GMV) significantly alters the observed BEN-task activation relationship.

In this study, we first replicated key prior findings, including the effects of sex and age on BEN and GMV, their spatial and brain-wide voxel-wise correlations, and the relationship between BEN and task activation across social cognition, reward, and emotion tasks. Subsequently, we examined the correlation between GMV and task activation, as well as the correlation between BEN and task activation after controlling for GMV.

Our results successfully replicated previous findings. Crucially, the relationship between BEN and task activation remained largely unchanged after controlling for GMV. This indicates that BEN captures unique variance in task activation beyond what is explained by GMV alone, establishing BEN as a functional bridge linking brain structure to task activation. Future work should employ predictive analyses to quantify the added value of BEN in predicting task activation from structural information.

## 1. Introduction

How the brain gives rise to diverse and complex functions from its fundamental structure to sustain our daily lives has been one of the most important questions in neuroscience. The development of multimodal neuroimaging and the accessibility of large-scale open datasets have continuously advanced this field. In this study, we utilize HCP multimodal neuroimaging dataset (Van Essen et al., 2013) along with brain entropy (BEN) (Wang, Li, Childress, & Detre, 2014), a powerful tool for measuring brain activity dynamics, to provide new insights into this ongoing inquiry.

BEN, derived from fMRI, has been widely validated as a reproducible and sensitive metric of brain activity (Del Mauro & Wang, 2024a; Lin, Chang, Song, Li, & Wang, 2022; Wang, 2021; Wang et al., 2014). In addition, recent studies have established links between resting-state BEN and both task-evoked brain activity and brain morphology. Specifically, lower resting-state BEN is brain-wide associated with heightened task activation and deactivation (Lin et al., 2022), exhibits brain-wide voxel-wise negative correlations with structural features such as gray matter volume (GMV) and cortical surface area (Del Mauro & Wang, 2024b) (Del Mauro & Wang, 2024b), while also demonstrating a spatial positive correlation with GMV (Song & Wang, 2024) (D. Song & Z. Wang, 2024).

However, the specific role of BEN in the relationship between brain structure and task activation, and whether it provides unique information, remains unclear. This study first replicates the established findings on the associations between BEN and task activation, as well as GMV. To confirm the specificity of the BEN-task activation relationship, we regressed out GMV information from both the BEN and task activation maps. This procedure allowed us to test whether the relationship was significantly altered and to determine the extent to which it is underpinned by structural morphology.

## 2. Methods

### 2.1 Dataset

All participants from the Human Connectome Project (HCP) 3T S1200 release (Van Essen et al., 2013). The final sample consisted of 649 participants with 4 resting-state runs, a T1-weighted structural image, and functional data for 3 distinct tasks, including social cognition, gambling, and emotion processing (Barch et al., 2013). All participants were healthy individuals between 22 and 35 years (mean age = 28.72, standard deviation = 3.68). The MR images were scanned on a 3T Siemens Skyra scanner using a standard 32-channel head coil at Washington University and preprocessed by the HCP team. Detailed information about data acquisition and preprocessing can be found in (Glasser, Sotiropoulos et al. 2013, Van Essen, Smith et al. 2013).

### 2.2 GMV

GMV was computed from “T1w_acpc_dc_restore_brain.nii.gz” by the FSL pipeline (Smith, Jenkinson et al. 2004, Douaud, Smith et al. 2007) following FSLVBM protocol (https://fsl.fmrib.ox.ac.uk/fsl/fslwiki/FSLVBM), fslvbm_2_template was used to create the study-specific GM template, fslvbm_3_proc was used to register all GM images to the study-specific template non-linearly. GMV maps were smoothed using an isotropic Gaussian kernel (full width at half maximum = 4 mm). The detailed guide of FSLVBM is available at https://fsl.fmrib.ox.ac.uk/fsl/fslwiki/FSLVBM/UserGuide.

### 2.3 BEN

BEN maps were calculated from preprocessed resting-state fMRI using the BEN mapping toolbox (BENtbx) (Wang et al., 2014) based on sample entropy (SampEn) (Richman & Moorman, 2000). SampEn is a measure designed to quantify irregularity of a time series by assessing the probability of similar patterns occurring within the sequence. It evaluates the likelihood that two sequences of length *m* and *m*+1 will remain similar under a given tolerance threshold *r*. In the study, the window length (dimension) was set to *m* = 3, and the cut-off threshold was set to *r* = 0.6 according to Wang et al (2014) experimental optimization after evaluating the impact of a range of window lengths (*m*) and threshold values (*r*) on the results (Wang et al., 2014). Following averaging BEN maps across the four runs, the BEN maps were smoothed using an isotropic Gaussian kernel (FWHM = 4 mm).

### 2.4 Task

This study utilized task fMRI data from three paradigms: social cognition, gambling, and emotion processing. The data, which were processed by the HCP team and the group-average activation maps can be accessed from ConnectomeDB (https://www.humanconnectome.org/study/hcp-young-adult/article/s1200-group-average-data-release). Our analysis used the task activation z-value maps from two runs across 649 individuals. For detailed experimental design, please refer to (Barch et al., 2013).

### 2.5 Statistical analysis

First, the independent effects of sex and age on BEN and GMV were evaluated separately, with each model mutually adjusted for the other variable (i.e., the effect of sex was controlled for age, and vice versa).

The average maps for BEN, GMV, and task activation z-values were first derived. Spatial correlations among these maps were then assessed using both raw and absolute values. The brain-wide voxel-wise correlation between BEN and GMV was evaluated after controlling for age and sex.

After controlling for sex and age, correlation analyses were conducted between both BEN and GMV and various task-activated z-value maps. Subsequently, the relationship between BEN and the task activation maps was re-examined after regressing out the effects of sex, age and GMV. The spatial correlations between these correlation maps and the average BEN, average GMV, and average task activation maps were also evaluated.

For the brain-wide voxel-wise correlation analysis, the significance threshold was set at *p* < 0.001 at the voxel level. Given the large number of voxels (*n* = 228,483), even minimal correlations could yield extremely small p-values. Therefore, for the spatial correlation analysis, we employed an empirical threshold of *r* > 0.1 to define a meaningful correlation, a level at which the corresponding p-values were far smaller than 10 ^-10^.

## 3. Results

### 3.1 Sex and age effects on BEN and GMV

As illustrated in Figure 1, lower levels of both BEN and GMV were observed in males compared to females. Furthermore, age was positively correlated with BEN but negatively correlated with GMV, replicating previous findings (Bethlehem et al., 2022; Wang, 2021).

**Figure 1.**
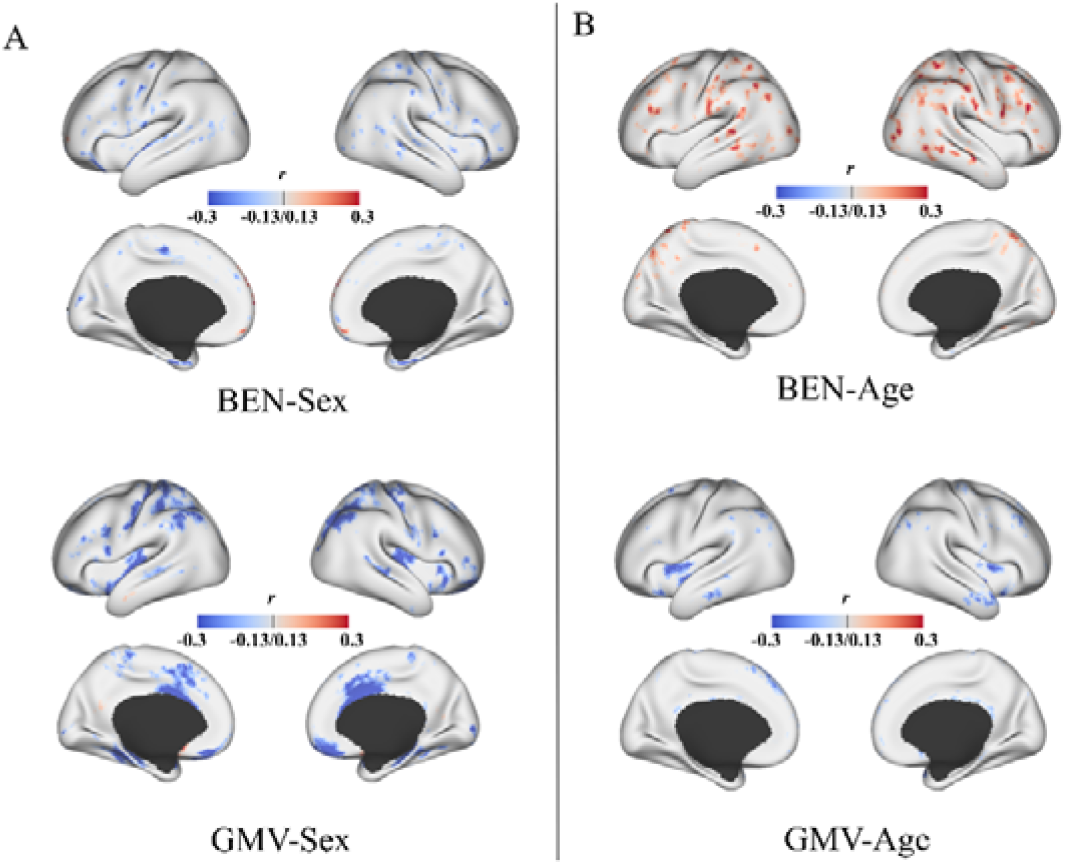
Sex and age effects on BEN and GMV. A. Sex effect on BEN and GMV, cool color indicates lower BEN or GMV in male. B. Age effect on BEN and GMV, warm color indicates positive correlation, and cool color indicates negative correlation.

### 3.2 Average BEN, GMV, and Task Activation

The average BEN, GMV, and task activation z-value maps are presented in Figure 2. Replicating prior findings, a brain-wide voxel-wise correlation analysis revealed a significant negative correlation between BEN and GMV (Del Mauro & Wang, 2024b; Lotze et al., 2019).

**Figure 2.**
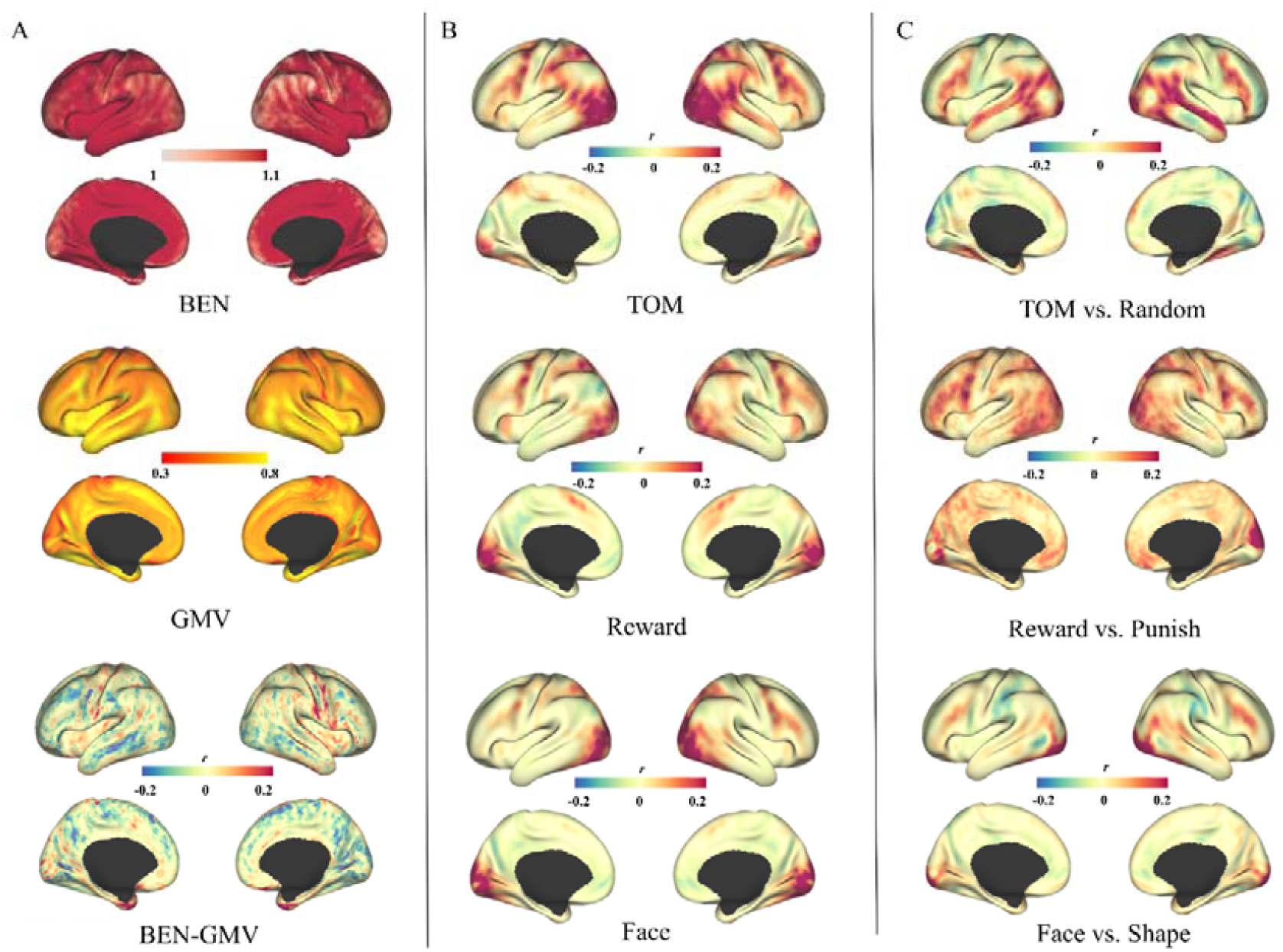
Average BEN, GMV, and Task Activation. A. From top to bottom: the average BEN map, the average GMV map, and the threshold-free correlation map between BEN and GMV. Warm color indicates positive correlation; cool color indicates negative correlation. B. Mean z-value maps for task-versus-baseline contrasts, displayed from top to bottom for the Theory of Mind (ToM), Reward, and Face tasks. C. Mean z-value maps for direct task contrasts, displayed from top to bottom as: ToM vs. Random, Reward vs. Punishment, and Face vs. Shape. Warm and cool color represent positive and negative activation, respectively.

At the spatial map level, BEN and GMV demonstrated a significant positive spatial correlation (*r* = 0.32, *n* = 228,483), consistent with previous results derived from 400 cortical parcels in the HCP 7T dataset (D. Song & Z. Wang, 2024). While BEN showed no significant spatial correlation with any of the task activation maps using either raw or absolute values, GMV was positively correlated with various task activations. Furthermore, the task activation maps were broadly positively correlated with one another. These correlations were further enhanced when absolute activation values were used. And, as illustrated in Figure 3, spatially regressing GMV out from the maps did not substantially alter the correlation patterns between BEN and task activation, or the inter-correlations among the task activations themselves.

**Figure 3.**
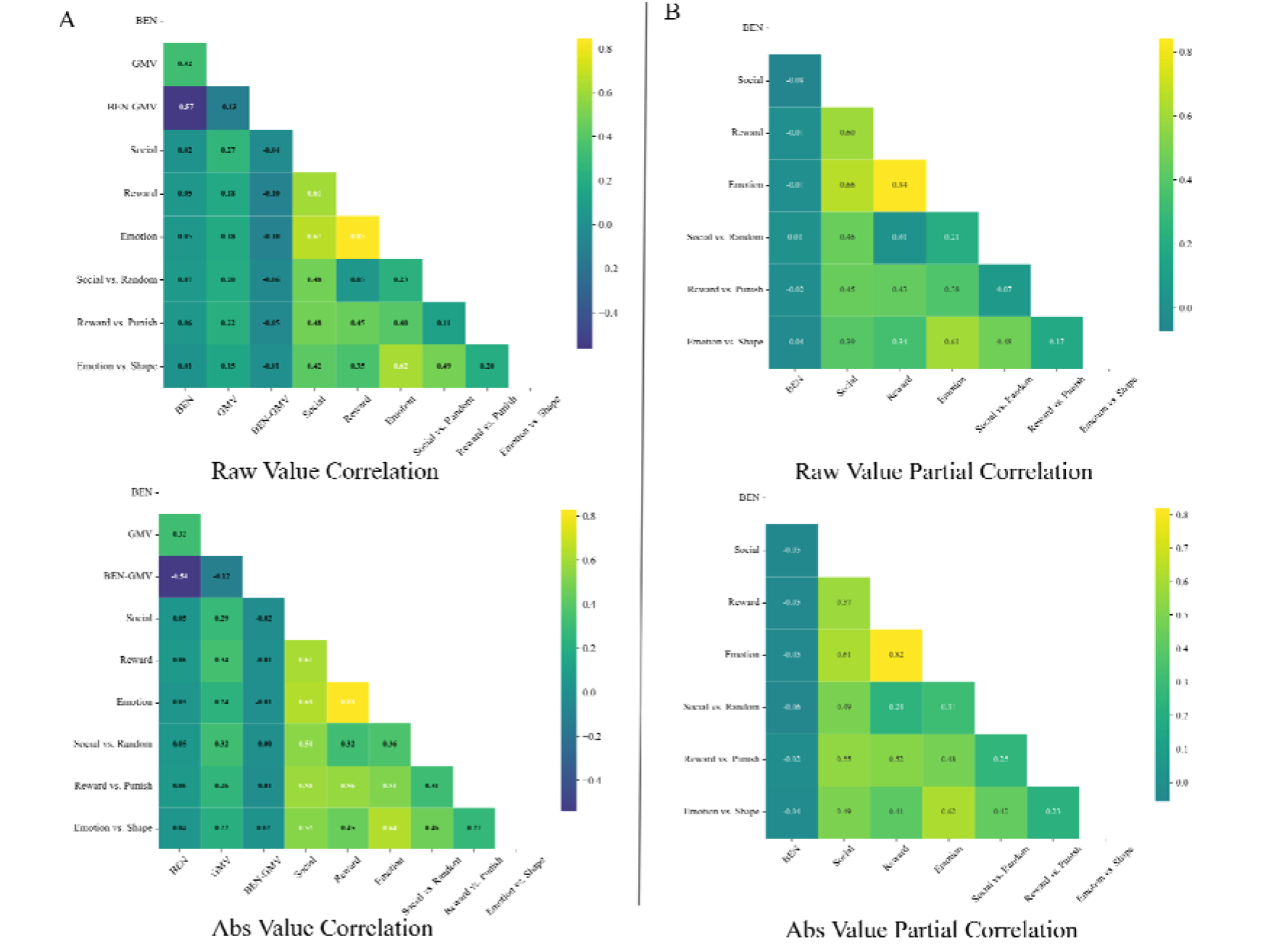
Spatial correlations among group-average maps. A. Correlations between the original average maps. Top: raw values; Bottom: absolute values. B. Correlations between residual maps after spatially regressing out GMV. Top: raw residuals; Bottom: absolute values of residuals. Warm/cool colors denote positive/negative correlations in all panels.

### 3.3 Correlations of BEN and GMV with Task Activation

BEN demonstrates a negative correlation with task vs. baseline activation, manifesting as stronger negative correlations in regions with higher positive activation and stronger positive correlations in regions with higher negative activation. Spatially, higher BEN values are observed in positively activated areas, while lower BEN values are found in negatively activated regions. Even after regressing out GMV, the correlation pattern between BEN and task vs. baseline activation remains consistent. This stability persists even when examining more complex inter-task activation contrasts. In contrast, GMV shows a positive correlation with task activation, where greater GMV is associated with higher levels of task-induced activation. These results are presented in Fig. 4 and Fig. 5.

**Figure 4.**
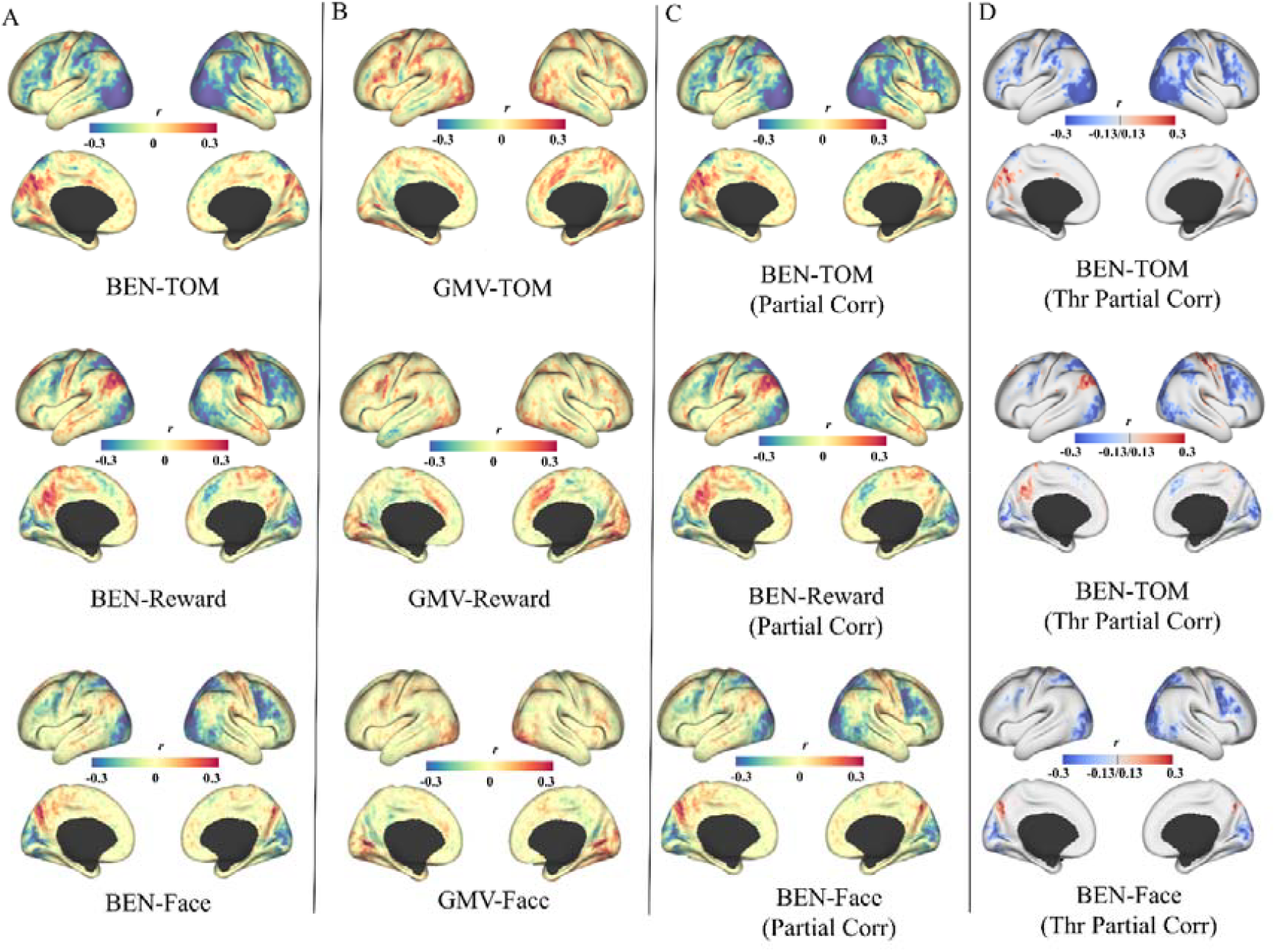
The correlations between BEN, GMV and task vs. baseline activation. A. BEN vs. task activation (controlling for sex and age). B. GMV vs. task activation (controlling for sex and age). C. Partial correlations of BEN with task activation (controlling for sex, age, and GMV). D. Thresholded map of the partial correlations in panel C. In all maps, the three rows correspond to the ToM, Reward, and Face tasks (top to bottom). The color scale denotes the correlation coefficient, with warm and cool colors representing positive and negative values, respectively.

**Figure 5.**
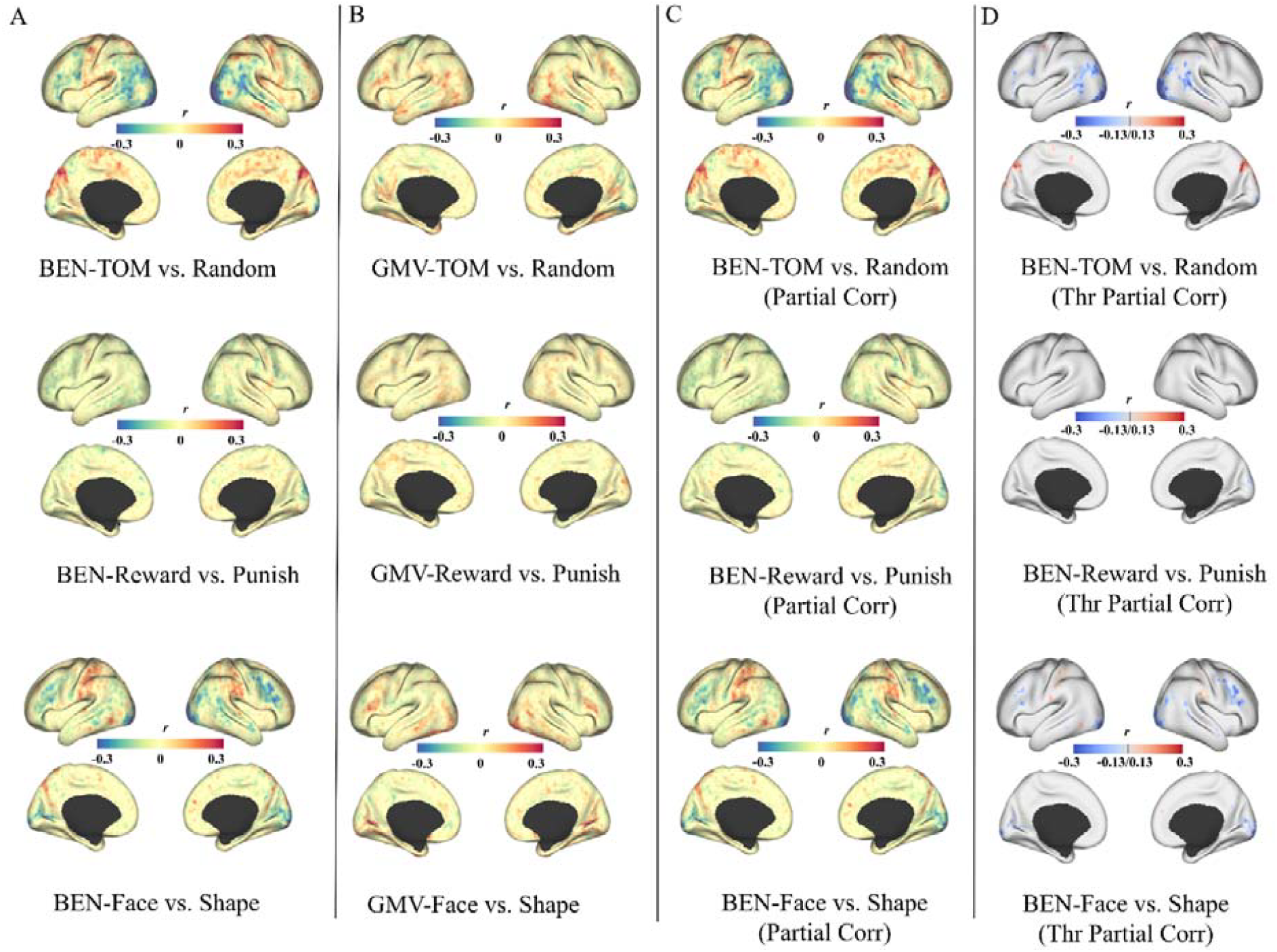
The correlations between BEN, GMV and task contrast activation. A. BEN vs. task activation (controlling for sex and age). B. GMV vs. task activation (controlling for sex and age). C. Partial correlations of BEN with task activation (controlling for sex, age, and GMV). D. Thresholded map of the partial correlations in panel C. In all maps, the three rows correspond to the ToM vs. Random, Reward vs. Punish, and Face vs. Shape tasks (top to bottom). The color scale denotes the correlation coefficient, with warm and cool colors representing positive and negative values, respectively.

**Figure 6.**
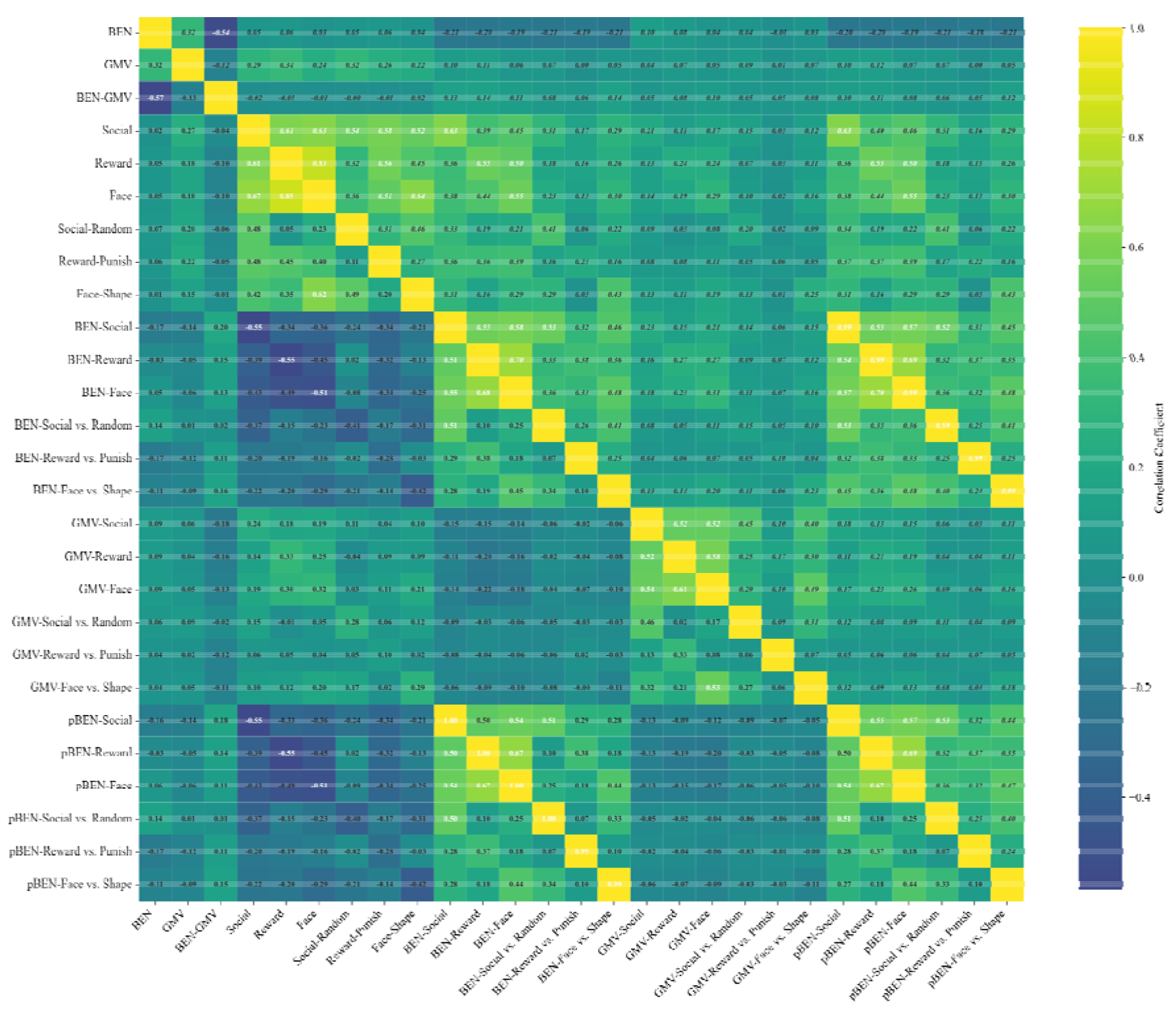
Spatial correlations among average BEN, GMV, task activation and correlation maps. The lower triangle of the matrix displays correlations based on the original values, while the upper triangle displays correlations based on the absolute values. The color intensity reflects the magnitude of the correlation, with warm and cool color scales representing positive and negative values, respectively.

Spatial correlation analysis further reveals that the spatial distribution of the correlation between BEN and task activation is markedly negatively correlated with the spatial pattern of mean task activation. This relationship remains largely unchanged regardless of whether GMV is regressed out (see Fig.6).

## 4. Discussion

This study demonstrates that BEN is not entirely determined by brain structure but provides substantial functional information, thereby building a bridge from brain structural architecture to the realization of complex brain functions. Furthermore, as a characteristic of local brain activity, BEN serves as a foundational element for the formation of brain networks. This paves the way for future research to understand how the brain progresses from structure to local function, and further to the establishment of brain networks, ultimately elucidating how complex cognitive functions emerge from their structural underpinnings.

### 4.1. BEN and GMV: Spatial Positive but Inter-individual Negative Correlation

Within individuals, BEN and GMV exhibit a positive spatial correlation in the study and our a previous study (D. Song & Z. Wang, 2024), whereas across individuals, they show a negative correlation in the study and a study from (Mauro & Wang, 2025). While this pattern may appear contradictory, it in fact reflects distinct aspects of brain organization. The spatial relationship between BEN and GMV reflects the structural basis of BEN. In the emergence of BEN from its structural underpinnings, neurotransmitters are likely to play a critical role. This is supported by our previous study, which found that neurotransmitters facilitate the structure-function coupling between BEN and GMV (D. Song & Z. Wang, 2024). Based on the interplay of inter-individual negative correlations and intra-individual positive spatial correlations, we contend that BEN cannot be viewed as a static property, but rather as a dynamic process of acquiring, processing, and compressing information. The within-individual positive correlation suggests that regions with higher BEN may be associated with greater external stimulation or information acquisition, whereas regions with lower BEN may relate to more intensive information processing and compression. This interpretation aligns well with the observed spatial distribution of BEN: higher values are found in unimodal sensorimotor cortex, and lower values in multimodal association cortex.

### 4.2. BEN, GMV and Task Activation

Our finding of a negative correlation between BEN and task activation is consistent with the results reported by Wang (Wang, 2021) and Lin et al (Lin et al., 2022). In the present study, we further accounted for the structural characteristic of GMV and demonstrated that the negative correlation between BEN and task activation persists even after regressing out GMV. This indicates that BEN captures information beyond what is represented by brain structure, thereby serving as a functional bridge linking anatomical features to the manifestation of complex task-induced activation.

We observed a positive spatial correlation between the average GMV map and the average task activation maps. Additionally, consistent positive spatial correlations were found among the task activation maps themselves. Notably, these inter-task spatial correlations persisted even after regressing out the spatial pattern of GMV. These findings suggest that task activation patterns share a common organizational basis, rooted in both a unified structural framework and a consistent functional architecture. In contrast, no significant spatial correlation was detected between the average BEN map and the average task activation maps, regardless of whether GMV was controlled for. This indicates that the spatial organization of task activation and that of BEN do not share the same functional principles. Therefore, BEN may represent a relatively distinct and independent feature of brain functional organization.

In this study, we employed spatial correlation analysis to determine whether the inter-subject correlation of BEN mirrors the spatial distribution of the average task activation pattern. Our results revealed a strong inverse similarity, a significant negative correlation between the voxel-wise inter-subject correlation of BEN and the spatial pattern of task activation. This inverse relationship stems from the negative correlation between BEN and task activation across individuals. Specifically, lower resting-state BEN was associated with higher levels of both task-positive activation, primarily in the lateral prefrontal and visual cortices and task-negative deactivation mainly within the default mode network. This pattern suggests that a lower BEN signifies a greater capacity for mobilizing computational resources and suppressing irrelevant activity, reflecting a state of enhanced information compression.

Furthermore, our previous research demonstrated that during movie-watching, BEN increases in the association cortices but decreases in the audiovisual regions compared to the resting state (D.-H. Song & Z. Wang, 2024). This profile that elevated BEN in higher-order areas alongside reduced BEN in sensory areas may indicate that complex task demands impose an information load that challenges processing capacity in associative regions, while simpler perceptual information is efficiently processed in sensory areas. The ability to compress information, reflected in the capacity to reduce BEN, appears to be linked to regional brain connectivity. This is consistent with our prior finding of a spatial negative correlation between regional BEN and whole-brain connectivity strength (Song, 2024).

### 4.3 Future Prospects

While the role of BEN as a bridge linking brain structure to complex task activation has emerged, several aspects of this initial manuscript require more detailed consideration and supplementation as the starting point of the research project. (1). Regarding sex and age differences, it would be appropriate to regress out either BEN or GMV. Given the negative correlation between GMV and BEN, and the fact that sex differences in GMV and BEN align in the same direction, more refined analyses may enhance the detection of these sex-related differences. (2). Statistical analyses should employ more appropriate thresholds. In this study, we used a voxel-level threshold of *p* < 0.001, while in spatial correlation analyses, we empirically adopted *r* > 0.1, primarily because permutation tests also yielded significant results even at very low *r*-values. A more objective standard for thresholding should be established in subsequent work. (3). We did not perform predictive analyses with cross-validation. Using BEN and GMV separately or in combination within a cross-validation framework would help quantify the extent to which BEN improves the prediction of task activation. Although we previously conducted a preliminary test with about 100 participants, suggesting that combining GMV with BEN significantly enhances predictive performance for task activation, these findings require further consolidation and validation and have thus not been included in the current manuscript. (4). All analyses included subcortical and cerebellar regions in this study, the corresponding results have not been presented in this manuscript due to the large volume of figures. Furthermore, more detailed examinations should be conducted regarding the sensorimotor-association (S-A) axis of cerebral cortex, cerebellum and subcortex. The S-A axis represents a major axis of hierarchical cortical organization and the influence of this axis on BEN has been demonstrated in prior studies (D.-H. Song & Z. Wang, 2024; Song, 2024).

## Acknowledgements

Data used in this study were provided by the Human Connectome Project, WU-Minn Consortium (Principal Investigators: David Van Essen and Kamil Ugurbil; 1U54MH091657), funded by the 16 NIH Institutes and Centers that support the NIH Blueprint for Neuroscience Research, and by the McDonnell Center for Systems Neuroscience at Washington University.

## Data and code availability

All MRI data are available at https://www.humanconnectome.org/.

BENtbx is available at https://www.cfn.upenn.edu/zewang/BENtbx.php.

## Notes

https://www.humanconnectome.org/

### Competing Interest Statement

The authors have declared no competing interest.

